# Transposable elements as new players to decipher sex differences in Parkinson Disease

**DOI:** 10.64898/2026.03.27.714370

**Authors:** Fernando Gordillo-González, Cristina Galiana-Roselló, Rubén Grillo-Risco, Irene Soler-Saéz, Marta R. Hidalgo, Haruhiko Siomi, Mie Kobayashi-Ishihara, Francisco García-García

## Abstract

We present a novel integrative analysis of transposable elements (TEs) in 4 single cell RNA-seq (scRNA-seq) datasets of postmortem substantia nigra pars compacta samples of Parkinson Disease (PD) patients matched healthy controls, with the objective of building a cell-type specific trustworthy atlas of TEs that may clarify the role of TEs in sex differences in PD. We have used the soloTE tool to evaluate the TEs expression changes across all snRNA-seq studies identified in our previous systematic review, and then integrated the results using meta-analysis techniques. Finally, we evaluated the possible associations between TEs and protein coding genes by integrating our previous results in this matter with the information of TEs obtained, in order to propose the possible action mechanism by which some of the TEs contribute to PD.

## Introduction

Parkinson’s disease (PD) is the second most prevalent neurodegenerative disease after Alzheimer’s, as well as the leading movement disorder. Current estimates indicate that approximately 11.77 million individuals are affected worldwide, with both the incidence and prevalence of the disease showing a significant upward trend in recent years, primarily due to aging populations (https://doi.org/10.3389/fnagi.2024.1498756). Neuropathologically, PD is primarily defined by the progressive loss of dopaminergic neurons within the substantia nigra pars compacta (SNpc). While the medulla and midbrain are predominantly affected in the initial phases, the pathology extends to cortical regions as the disease advances. This neuronal degeneration is responsible for the emergence of a wide range of both motor and non-motor symptoms, including bradykinesia, tremor, depression or dementia among others. Although this disease imposes a significant social and economic burden, no reliable biomarkers to provide a definitive diagnosis nor therapeutic intervention capable of halting the progression of PD is still available, making its clinical management a considerable challenge.

The causes of PD have not yet been fully understood, although several disease hallmarks have been described throughout years, including the presence of lewy bodies, failure of protein clearance pathways, oxidative stress, excitotoxicity and neuroinflammation. Furthermore, several risk factors predisposing individuals to the disease development have been identified, with age, pesticide exposure, certain genetic variants and sex among the most relevant.

Numerous studies highlight the significant impact of biological sex on various aspects of PD, such as pathophysiology, symptoms, and treatment response. PD exhibits a higher incidence and prevalence in males, with an earlier age of onset compared to females. Regarding symptoms, males tend to experience rigidity, freezing of gait, cognitive impairment, and urinary dysfunction, whereas the predominant symptoms in females are tremor, pain, depression or dyskinesias, among others. At the pathophysiologic level, female patients show higher striatal dopaminergic activity and lower oxidative stress and microglial activation, which may benefit from the neuroprotective and anti-inflammatory effects of estrogen, among others biological, genetic, hormonal or environmental factors. These sex-specific variations in PD underscore the importance of considering sex as a factor in research, and support the notion that there are molecular mechanisms that may contribute differently to PD progression in a sex-dependent manner, despite they are not yet completely understood.

To address this problem, high-throughput technologies, such as single cell RNA-seq (scRNA-seq), provide us with more holistic descriptions of the diseases and much deeper understanding of cellular heterogeneity within tissues. We recently conducted a sex-specific meta-analysis of scRNA-seq studies in postmortem brain tissue from PD patients, where we aimed to characterize sex differences in PD at both gene expression, cell-cell communication and pathway activation. From these results, it is remarkable the greater neuroinflammation pattern in males and, in terms of neurodegeneration, it was more pronounced in the SNpc of males but, regarding the cortex, were more prevalent in females (https://doi.org/10.1101/2024.12.20.628852). However, in this study we only took into account the protein-coding genes, not the non-coding fraction of the genome, whose characterization has been the focus of recent scientific efforts.

In recent years, Transposable Elements (TEs) are emerging as new players in the study of neurodegenerative diseases, as they constitute a major part of the human genome (∼40-50%) and have been recognized to play important roles in genomic stability, gene expression, and cellular function through alterations in their expression (PMCID: PMC9186530). Although the existence of several silencing mechanisms of TEs to maintain the genome integrity, it is described that during aging there is a triggering of TE activation, mainly caused by the disruption of heterochromatin structure and the loss of silencing proteins (Ravel-Godreuil et al., 2021). This TE dysregulation has been related to genomic instability, apoptosis and inflammation, emerging, therefore, as a major hypothesis to explain aging-associated diseases such as Parkinson’s disease. In this context, computational pipelines and tools have emerged to quantify TEs expression levels from transcriptomics data, since it has been proposed that aberrant transcription of TEs may be associated with age-related disease processes. For bulk RNA-Seq, several tools have been developed to analyze TE expression (PMID: <u>26206304</u>, PMID: <u>35876845</u>, PMID: <u>40140852</u>), however, such software fails to detect specific expression patterns of cell populations. Recently, Rodríguez-Quiroz et al. developed SoloTE, a pipeline for the single cell analysis of TE expression, allowing the study of these poorly characterized players on a cell-by-cell basis.

Various studies have related the presence of certain polymorphisms of TEs (PMCID: PMC6102640, PMCID: PMC4856358) and the increase their expression to the PD pathology (PMCID: <u>PMC9597212</u>, PMCID: <u>PMC11526979</u>), highlighting the particular involvement of LINE-1 (L1) and ERV families, although the processes by which TEs may contribute to the neurodegenerative process are not fully dissected. Additionally, this evidence has been largely supported by disease models or bulk technologies in postmortem brain tissue, masking the possible TE cell-specific transcription dynamics (Mustafin, 2025). Few recent studies have addressed this pitfall by applying snRNA-seq technology to explore these activity differences, describing cell-type specific TE patterns that might be related to hallmark PD processes, such as neuroinflammation (Deng et al., 2024; Garza et al., 2025). However, some authors have also described a dynamic and inconsistent TE expression across snRNA-seq datasets (Deng et al., 2024), making this field a suitable candidate for integrative strategies such as a meta-analysis. Yet, to our knowledge, no robust meta-analysis approach that ensures the current findings are not dependent on a specific dataset have been applied to date. Also, sex differences in the deregulation of TEs remains largely unexplored, as none of studies examining TEs in PD performs sex-stratified analyses, despite the fact that population-based studies have identified sex differences in TE expression, possibly related to Y chromosome TE content (<u>10.1101/2023.08.03.550779</u>, Talevi et al., 2025). The complete understanding, at a cellular level, of these sex-differences and their related biological mechanisms may open new avenues for therapeutic interventions aimed at modulating their activity or mitigating their detrimental effects during PD in a sex-specific manner.

In this work, we present a novel integrative analysis of TEs in 4 snRNA-seq datasets of postmortem SNpc samples of PD patients matched healthy controls, with the objective of building a cell-type specific trustworthy atlas of TEs that may clarify the role of TEs in sex differences in PD. We have used the soloTE tool to evaluate the TEs expression changes across all snRNA-seq studies identified in our previous systematic review, and then integrated the results using meta-analysis techniques. Finally, we evaluated the possible associations between TEs and protein coding genes by integrating our previous results in this matter with the information of TEs obtained, in order to propose the possible action mechanism by which some of the TEs contribute to PD.

## Methods

### Systematic Review

We performed a systematic review in accordance with the Preferred Reporting Items for Systematic Reviews and Meta-Analyses (PRISMA) guidelines, which emphasize standardization and reproducibility. Initially, public databases, including GEO (Gene Expression Omnibus), ENCODE, UCSC Cell Browser, Human Cell Atlas, and Single Cell Expression Atlas, were queried using the search terms: “Parkinson’s,” “Parkinson’s disease,” or “PD.” To ensure the relevance of the datasets, we applied the following inclusion criteria: i) organism: Homo sapiens, and ii) the study type had to involve single-cell transcriptomics (scRNA-seq or snRNA-seq). Studies were excluded if they met any of the following exclusion criteria: i) experimental design not including healthy controls, ii) the research focus was not on PD, iii) the samples were not derived from post-mortem brain tissue, iv) lack of information on patient sex, v) lack of representation of both sexes in all conditions, or vi) not accessible raw data. Subsequently, raw data from the eligible studies were retrieved from the SRA repository using the SRAToolkit package. Results from PRISMA analysis performed in our previous work can be consulted in Supplementary Figure 1 (Figure S1).

### Alignment and TE count inference

Once we have the fastq files with the original reads for each sample, we perform a quality control using fastqc (v0.11.8) and multiqc (v1.12). After verifying the integrity of the data, we use Cellranger (v7.1.0) to align the reads with the human GRCh38 pre-mRNA reference genome, with the default parameters and providing multiple input fastq files for multi-lane libraries. Next, we used the bam files obtained from the alignment to infer the transcriptional activation of the TEs with the soloTE package (v1.09), using the annotation file provided in the tool and the default parameters.

### Individual snRNA-seq data processing

We first set a SingleCellExperiment (SCE) object that contained all the information from the soloTE output, using SingleCellExperiment (Amezquita et al., 2020) and DropletUtils (Lun et al., 2019) R packages. For quality control, we used the thresholds from the original paper for the number of Unique Molecular Identifiers (UMIs) and detected genes, along with a percentage of mitochondrial genes higher than 5%, to discard empty droplets or damaged cells. Genes with less than 5 counts were also removed. Finally, we discarded those droplets considered as doublets by the scDblFinder R package, as well as those with a library size larger than 3 MADs (Mean Absolute Deviation). After filtering the cells, we normalized the data with scran (Lun et al., 2016) and scater R packages.

We also employed scran and scater packages to select the 25% the most variable genes and perform a principal component analysis (PCA) for dimensionality reduction, using PCAtools R package (Blighe & Lun, 2021) to select the number of principal components (PCs) that summarize the largest amount of variance. With these PCs we performed and plotted the Uniform Manifold Approximation and Projection (UMAP) for visualization purposes. For cell clustering, the scran package was applied, combining the k-means algorithm with the Shared Nearest Neighbors (SNN) approach. The number of centroids and neighbors was adjusted based on the cell count in each study to ensure consistent resolution across datasets. Cell annotation was transferred from the cell types found in our previous gene expression atlas (<u>10.1101/2024.12.20.628852</u>). For identifying cell-type marker TEs, we converted the SCE object to SeuratObject and used the FindAllMarkers function from the Seurat package. The results were filtered for TEs with an FDR <0.05 and a logFC > 1.

### TE Differential expression analysis

Differential expression analyses (DEA) of TEs were conducted using the MAST R package (McDavid et al., 2021). A hurdle model was constructed using the *zlm* function, where the main explanatory variable was the combination of disease condition and sex (Control.females, PD.females, Control.males, PD.males). To account for potential intra-patient variability and avoid treating nuclei from the same donor as independent observations, the sample origin of each nucleus was included as a covariate in the model. Statistical significance was assessed using the lrTest function, while the getLogFC function was used to derive the logarithm of fold change (logFC) and the standard deviation (SD) for each TE. The p-value was corrected using the Benjamini & Hochberg (BH) algorithm.

To characterize TE expression changes in PD while accounting for sex differences, four contrasts were evaluated, three pairwise and one double contrast. The 3 pairwise comparisons included: i) PD vs Control (PDvsC) comparison, that captures disease-associated changes regardless of sex; ii) PD.female vs Control.female, that represents the Impact of Disease in Females (IDF); iii) PD.male vs Control.male, that represents the Impact of Disease in Males (IDM). These contrasts compare two experimental groups, with the group following “vs” used as the reference. Therefore, a positive logFC indicates increased expression in the case group relative to the reference, whereas a negative logFC indicates decreased expression.

In addition, we evaluated a sex-by-disease association contrast, referred to as the Sex Differential Impact of Disease (SDID), designed to identify TEs whose PD-associated expression changes differ between females and males. This contrast was defined as:

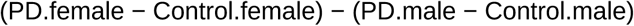

In this contrast, a positive logFC indicates that the magnitude of the PD-associated expression change is greater in females than in males, regardless of whether the TE is upregulated or downregulated. Conversely, a negative logFC indicates a greater disease-associated change in males. A schematic representation of the possible scenarios for this contrast is provided in Supplementary Figure 2 (Figure S2).

### Meta-analysis approach

We used all the individual results of all the studies available to perform a meta-analysis at the TE level for each cell type identified. The meta-analyses were conducted utilizing the R package metafor (Viechtbauer, 2010), employing the DerSimonian and Laird random effects model to account for heterogeneity across individual studies. This approach assigns greater weight to studies with lower variability in their results. For each feature (TE or gene), the analysis computed key statistical metrics, including the p-value, log fold change (logFC), and standard deviation (SD), among others. To control for multiple testing, the p-values were adjusted using the Benjamini & Hochberg (BH) correction method considering a feature with an adjusted p-value < 0.05 as significantly differentially expressed TE (DETE) or gene (DEG). To ensure the robustness of the findings and mitigate potential biases, diagnostic tests were applied to the results. These included the generation of funnel and forest plots, as well as influence and sensitivity analyses, such as leave-one-out cross-validation, to assess the stability and reliability of the outcomes.

### TE-Gene association network

To integrate the differential expression results of coding genes and TEs, we used the logFC values obtained from all contrasts (IDF, IDM, SDID, and PDvsC). We calculated Pearson correlations between DETEs and DEGs to identify TE–gene associations from a sex-specific perspective. Only correlations with an associated p-value < 0.01 and absolute correlation values > 0.9 were retained for downstream analyses. These significant TE–gene pairs, together with the SDID logFC values, were then used to construct TE–gene association networks in Cytoscape (v3.10.2).

To provide additional support for the predicted associations, we performed a genomic distance analysis between each TE–gene pair. The distance for each association was defined as the distance between the closest TE locus and the transcription start (if TE is located downstream) or end (if located upstream) site of the gene (See Methods 8). Associations with genomic distances < 10 kilobases, as well as intragenic TE insertions, were flagged and prioritized within the networks.

### TE-Gene distance analysis and contrasts

Genomic coordinates TEs were obtained from the BED annotation provided with soloTE, ensuring consistency with the annotation used for TE quantification. Because each TE subfamily comprises multiple genomic insertions distributed across the genome, TE-gene distances were defined using the nearest TE locus for each TE-gene pair. Distances were calculated relative to the gene transcription start site (TSS) when the TE was located downstream of the gene, and relative to the gene end coordinate when the TE was located upstream. TE insertions overlapping the gene body were classified as intragenic and assigned a distance of 0 base pairs.

To generate the distance datasets used in the comparisons, all TEs and protein-coding genes of interest (e.g., DETEs and DEGs identified in neurons) were iteratively paired, and the minimum genomic distance between each TE-gene pair was computed. For each pair, the distance in base pairs and the relative genomic position of the TE with respect to the gene (upstream, intragenic, or downstream) were recorded.

Differences between distance distributions were assessed using a one-tailed Mann-Whitney-Wilcoxon (MWW) nonparametric test, comparing distances between correlated versus non-correlated DETE-DEG pairs. When multiple comparisons were performed, p-values were adjusted using the Benjamini–Hochberg correction. The effect size (r) was calculated to quantify the magnitude of the difference between distance distributions using the standardized statistic from the Mann-Whitney-Wilcoxon test, according to the following formula:

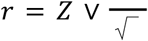

where Z is the standardized Mann-Whitney-Wilcoxon test statistic and N is the total number of observations used in the comparison.

### Functional enrichment

To identify biological programs associated with TE-linked gene regulation, we performed a biological characterization of the coding-gene sets included in the TE-gene association networks, for each cell type. The weight01 algorithm (Alexa et al., 2006) from the topGO R package (Alexa & Rahnenfuhrer, 2023) was used for the Biological Process (BP) GO ontology with a minimum GO term size of 15 genes. We defined as significant GO-BP terms, those with a p-value < 0.05. Graphical representations of the results were performed using ggplot2 and cnetplot function from clusterProfiler R package.

### Web platform development

We developed a user-friendly online resource to easily explore all the results obtained from the different analyses conducted in this work. The web app (https://bioinfo.cipf.es/cbl-atlas-pd/) was developed under the structure of the R shiny package (Chang W, 2024) and is compartmentalized into the different steps performed in the analysis.

## Results

### Individual analysis and TE expression

This study builds upon the datasets identified in our previous systematic review and meta-analysis of gene expression in PD. Raw sequencing data from these studies were retrieved and reprocessed to quantify TE expression. One cortex dataset was excluded due to incomplete raw files required for alignment. Consequently, the analysis was restricted to the four studies profiling the substantia nigra pars compacta (SNpc). The number of samples per experimental group across these datasets is shown in Figure 1A. Analysis with the soloTE package allowed us to identify transcripts associated with TE, representing, in most cases, around 10-20% of the total UMIs identified (Figure 1B). Although the average for all studies remained within this range, the GSE178265 study showed a greater dispersion of this percentage of TEs, reaching up to 40% in some of the samples analyzed.

**Figure 1.**
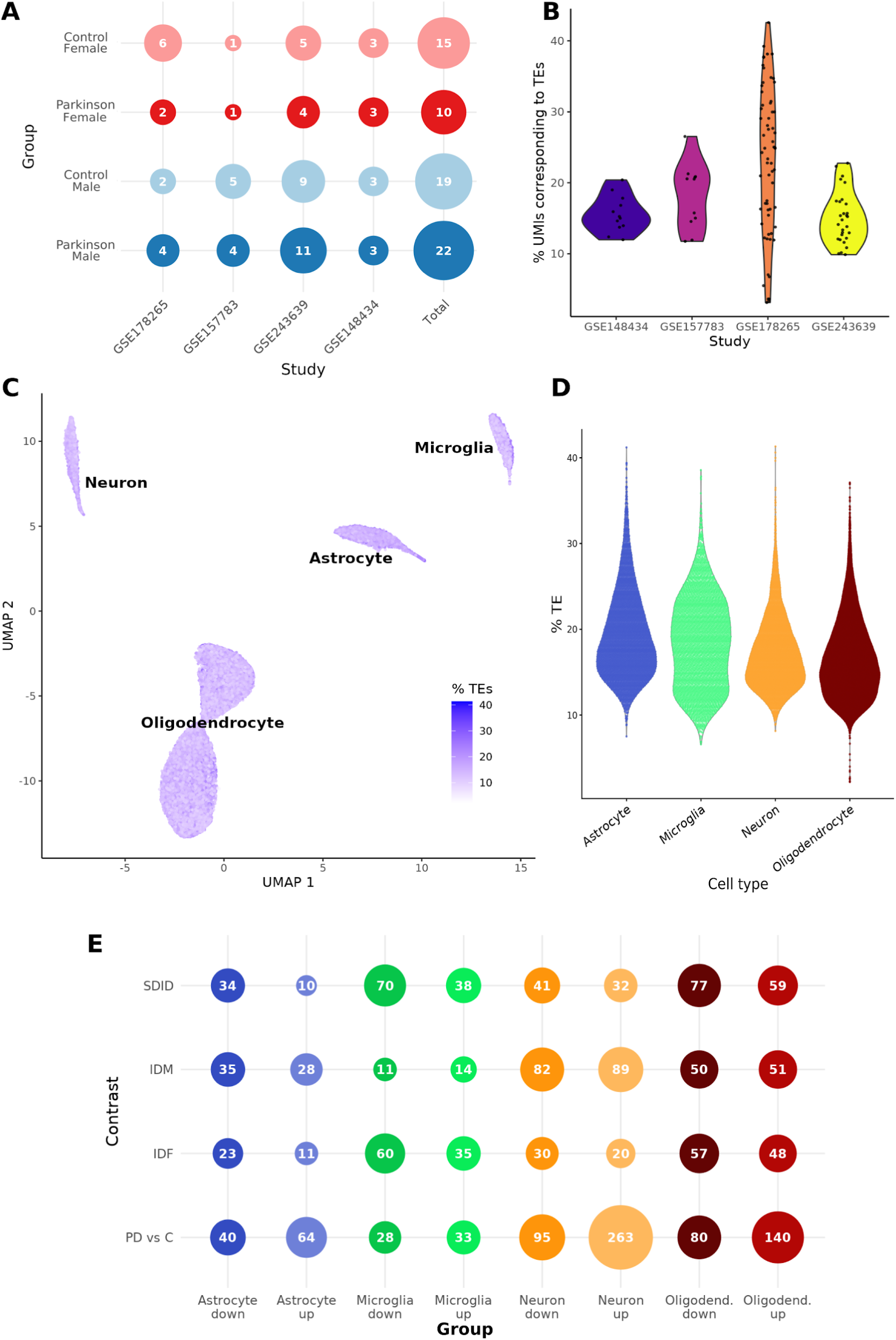
Individual analyses results overview and TE expression exploration. **A**-Number of individuals from each selected study, grouped by sex-disease condition. **B**-Violin plot showing the proportion of UMIs corresponding to TEs in each sample included in the meta-analysis. **C**-UMAP embedding of GSE157783 study, colored by the proportion of counts corresponding to TE transcripts. **D-** Violin plot of the TE transcript ratio per cell type of GSE157783 study. **E-** Number of significant DETEs of differential expression analyses performed in GSE157783 study, classified by contrast, cell type and logFC direction. TE: Transposable Element, DETE: differentially expressed transposable element, UMI: Unique Molecular Identifier, UMAP: Uniform Manifold Approximation and Projection, SDID: Sex Differential Impact of the Disease, IDM: Impact of the Disease in Males, IDF: Impact of the Disease in Females, PD vs C: Parkinson versus Control.

To further explore TE expression levels at single-cell resolution, we evaluated the proportion of TE-associated UMIs per cell across the different cell types previously identified. (reference paper 1). We selected the GSE157783 dataset as an example and performed a UMAP and violin plot to explore the distribution of TE expression by cell (Figure 1C-D). While TE expression levels exhibited some variability across different datasets, the overall landscape remained relatively consistent. Glial cells generally showed slightly higher TE expression compared to neurons, with most populations maintaining a median TE expression rate between 15-30% (Figure S3). Although certain neuronal clusters in specific cohorts showed lower proportions (under 10%), our integrated analysis confirms a widespread and significant TE presence across major brain cell types, regardless of the individual study lot.

The analysis of expression markers identified significant TE markers for all major cell types in all datasets included. However, both the expression levels and the proportion of cells expressing these TEs were generally lower than those typically observed for protein-coding markers, limiting their reliability as standalone cell-type markers (Figure S4A). Nevertheless, TEs displayed distinct cell-type–specific expression patterns that contribute to characteristic TE expression profiles across cell populations. One example is L1PA12, a L1 TE identified as a significant neuronal marker, expressed at a much higher fraction of cells (∼ 90%) than in the other glial cell types (∼ 30%) (Figure S4B). Another example is LTR46, a member of the ERV/LTR family, which was identified as a marker of oligodendrocytes with a high logFC (2.49). Despite the high logFC, UMAP visualization indicates that LTR46 is expressed only in a small fraction of cells (∼ 30%), but with a higher expression rate than the other cell types (Figure S4C). Interestingly, a comparative analysis of these TE markers across datasets revealed a small set of common TE markers for each cell type analyzed (8 in oligodendrocyte, 9 in astrocyte, 6 in neuron and 4 in microglia), constituting a consistent TE signal for each cell type that could collectively serve as a signature marker for the major brain cell types (Figure S4). Among these genes, the L1P4e subfamily in neurons is particularly noteworthy, as the L1 family has previously been reported to be more strongly expressed in this cell type. Similarly, the LTR9A1 and LTR12E subfamilies of the HERV family in oligodendrocytes are consistent with other single-cell studies (Deng et al., 2024).

Lastly, differential expression analyses at the TE subfamily level identified dysregulated DETEs in both positive and negative logFC across all cell types and contrasts performed (Figure 1E). Notably, neurons yielded the largest number of DETEs in most contrasts performed, with the exception of IDF, in which microglia achieved this qualification. Remarkably, neurons obtained the lowest number of DETEs in IDF which, in concordance with the results of SDID contrast that shows almost all the DETEs increased in males (logFC < 0), suggests less implication of TEs in this cell type in females in this study.

The number of DETEs identified in the remaining individual analyses can be found in the supplementary material (Figure S6), along with the web-based tool developed for data exploration. The number of DETEs varied considerably between studies (e.g., dataset GSE148434 showed the lowest number of DETEs in neurons but the highest in oligodendrocytes), highlighting the importance of statistically integrating evidence across studies rather than relying solely on results that are significant within individual datasets.

### Consensus signature of differential expressed TEs in PD

Following individual differential expression analyses, specific cell-type meta-analyses were performed on the results obtained in the four SNpc datasets so as to obtain robust results supported across studies. These meta-analyses were performed for all contrasts and the results were divided based on their logFC between up (>0) and down (<0) (Table 1). In the case of pairwise contrasts (IDM, IDF, and PD vs. C), the up category would represent higher expression in PD patients and vice versa for down, while in the case of the SDID contrast, up would imply an increase in the difference between women and down an increase in men (see Methods). By dividing the results between genes and TEs, we obtained a total of 537 DETEs (182 IDF, 81 IDM, 30 SDID, 244 PD vs C) and 14,457 DEGs (4,934 IDF, 3,981 IDM, 720 SDID, 4,822 PD vs C). The large difference between the results obtained for TEs and genes may be due to the difference in the number of starting elements for each MA (∼20,000 genes and ∼1,000 TE subfamilies), together with the lower stability in the expression of the former.

**Table 1.** Number of significative elements (FDR < 0.05) from meta-analyses in each cell type and contrast performed. TE: Transposable Element, SDID: Sex Differential Impact of the Disease, IDM: Impact of the Disease in Males, IDF: Impact of the Disease in Females, PD vs C: Parkinson versus Control.

|  |  | IDF |  | IDM |  | SDID |  | PD vs C |  |
| --- | --- | --- | --- | --- | --- | --- | --- | --- | --- |
|  |  | up | down | up | down | up | down | up | down |
| TEs | Astrocyte | 7 | 3 | 4 | 3 | 1 | 3 | 34 | 5 |
|  | Microglia | 5 | 3 | 8 | 1 | 0 | 0 | 3 | 2 |
|  | Neuron | 32 | 4 | 18 | 2 | 2 | 1 | 110 | 8 |
|  | Oligodendrocyte | 82 | 46 | 25 | 20 | 12 | 11 | 70 | 12 |
| Genes | Astrocyte | 923 | 394 | 792 | 328 | 43 | 96 | 776 | 152 |
|  | Microglia | 916 | 275 | 805 | 241 | 78 | 45 | 740 | 103 |
|  | Neuron | 271 | 370 | 248 | 202 | 83 | 122 | 403 | 369 |
|  | Oligodendrocyte | 914 | 871 | 859 | 506 | 140 | 113 | 1629 | 650 |

Comparative analyses of DETEs across contrasts and cell types revealed limited overlap between TE sets, suggesting that TE dysregulation is largely contrast- and cell-type-specific (Figure S7A-B). Across contrasts, oligodendrocytes showed the highest number of overlaps, all occurring between directionally consistent sets (e.g., 11 DETEs shared between PD vs C up and IDM up). A similar analysis performed for protein-coding genes identified a larger number of overlaps; however, this likely reflects the substantially higher number of DEGs compared with DETEs, as the largest gene sets remained contrast-specific. When overlaps were examined across cell types within the same contrast, only a few DETEs were shared between populations, further supporting the cell-type specificity of TE dysregulation. The PD vs C contrast showed the highest degree of overlap across cell types, highlighting TEs such as the FLAM-C subfamily, which was consistently downregulated in all four cell types analyzed. Most shared DETEs displayed concordant logFC directions, although a few showed opposite patterns, such as L1PA8A, which was upregulated in neurons but downregulated in astrocytes.

Previous studies have highlighted the contribution of LINE-1 (L1) and HERV families to PD (Mustafin, 2025), prompting us to examine their representation among the DETEs identified in our meta-analysis. Within the L1 family, the PD vs C contrast revealed dysregulated subfamilies across all cell types. Notably, neurons showed the strongest signal, with 39 L1 subfamilies significantly upregulated in PD (Figure 2A), with some subfamilies such as L1MC5 showing high consistency across datasets (Figure 2B-C), suggesting that L1 dysregulation is particularly pronounced in this cell population. Although relatively few sex-specific signals were detected, some L1 elements showed sex-dependent effects, such as L1M3c and L1MDa, which were higher in males than females in oligodendrocytes. Interestingly, nine L1 DETEs were identified in the IDF contrast compared with only one in IDM, which may indicate a stronger disease-associated dysregulation of L1 elements in females for this cell type (Figure 2A). For the HERV family (including ERV1, ERVK, and ERVL), the highest number of deregulated subfamilies was observed in oligodendrocytes, with 31 DETEs showing both up- and downregulation in the PD vs C contrast. In contrast to L1, HERV elements showed a clearer association with sex-related differences, being the most frequently deregulated TEs in the SDID contrast. Most of these elements were increased in males, with subfamilies such as LTR12C and Harlequin-int displaying particularly high logFC values (Figure 2D-E).

**Figure 2.**
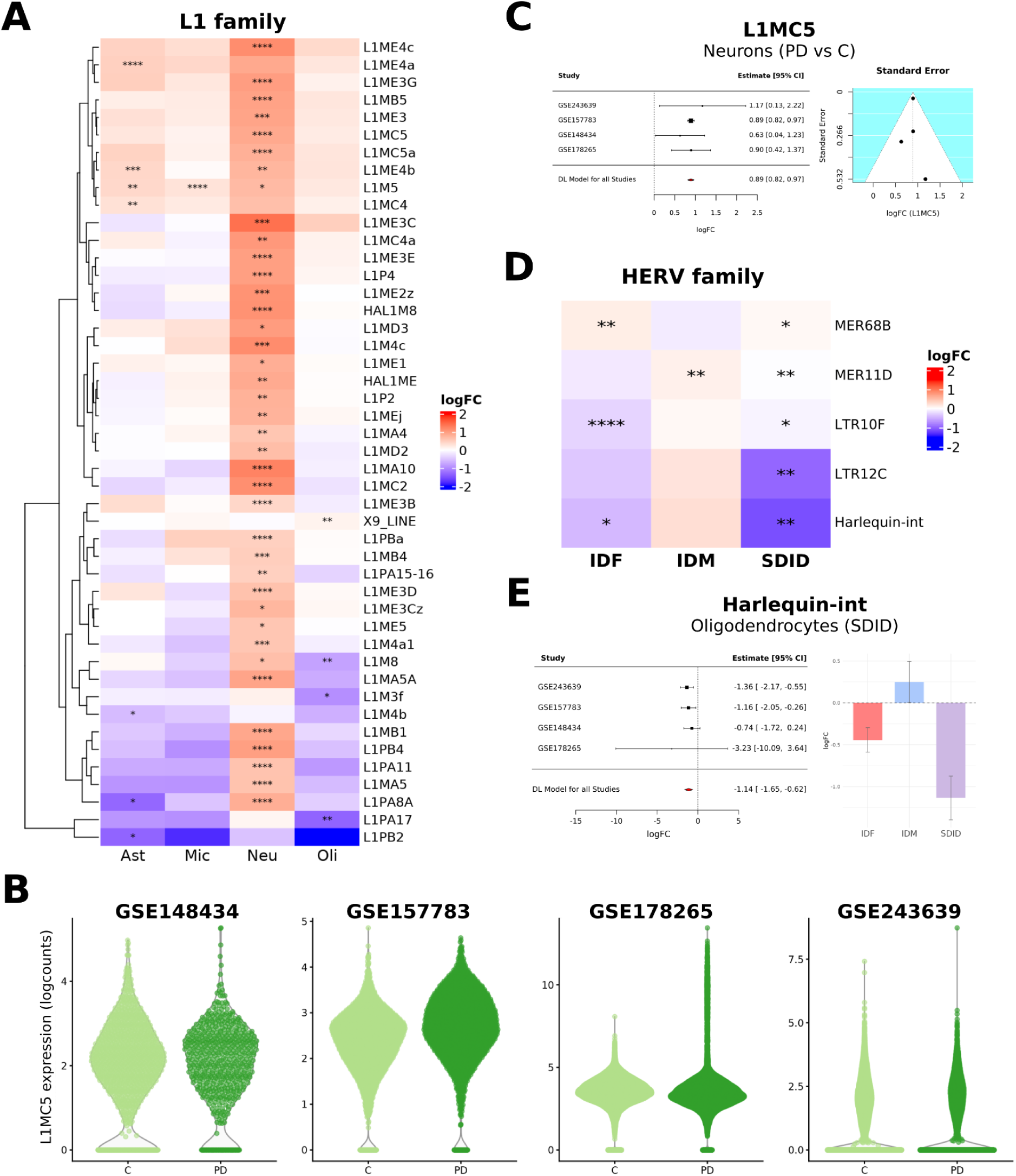
Meta-analysis results of relevant TEs. **A**-Heatmap of subfamilies of the L1 family that are significant in the PD vs C contrast, divided by cell type (*: FDR < 0.05, **: FDR < 0.01, ***: FDR < 0.001). **B**-Violin plot of expression distribution of L1MC5 subfamily in neurons per dataset, divided by condition. **C**-Forest (left) and funnel (right) plot of L1MC5 subfamily meta-analysis results in PD vs C contrast. **D**-Heatmap of HERV subfamilies that are significant in the SDID contrast in oligodendrocytes, divided by sex contrast. **E**-Forest plot (left) and barplot (right) of Harlequin-int subfamily meta-analysis results in SDID contrast. C: Control, PD: Parkinson’s disease, Ast: Astrocytes, Mic: Microglia, Neu: Neurons, Oli: Oligodendrocytes.

All results are made publicly accessible through a custom interactive web resource (available at https://bioinfo.cipf.es/cbl-atlas-pd/). This platform provides full access to the datasets and enables users to explore the data dynamically and generate customized plots.

### Biological characterization of TEs

The above results clearly demonstrate a novel link between TE expression and sex differences in PD. To further explore the nature of this association, we computed correlations between logFC values obtained across contrasts for both DETEs and DEGs. Significant correlations (p < 0.01) were interpreted as potential functional associations between TEs and genes, considering that concordant or anticorrelated regulation patterns may reflect relevant biological relationships.

To further characterize the relevance of these relationships, we examined the genomic distance between each TE locus and the correlated genes. Following the hypothesis that a shorter genomic distance has a higher probability of TE-gene association, we tested whether the distances between correlated DEGs and DETEs are shorter than those between non-correlated. We selected the closest locations of the DETE subfamilies to the obtained DEGs and grouped the distances according to correlated and uncorrelated DETEs, and then compared them using a MWW test. In microglia and neurons, the distances between correlated DETEs are significantly shorter (Figure 3A, S4D), while in oligodendrocytes, although not significant, results also suggest a shorter distance for correlated DETEs (p = 0.09) (Figure S4E). Astrocytes did not obtain significant results (Figure S4F).

**Figure 3.**
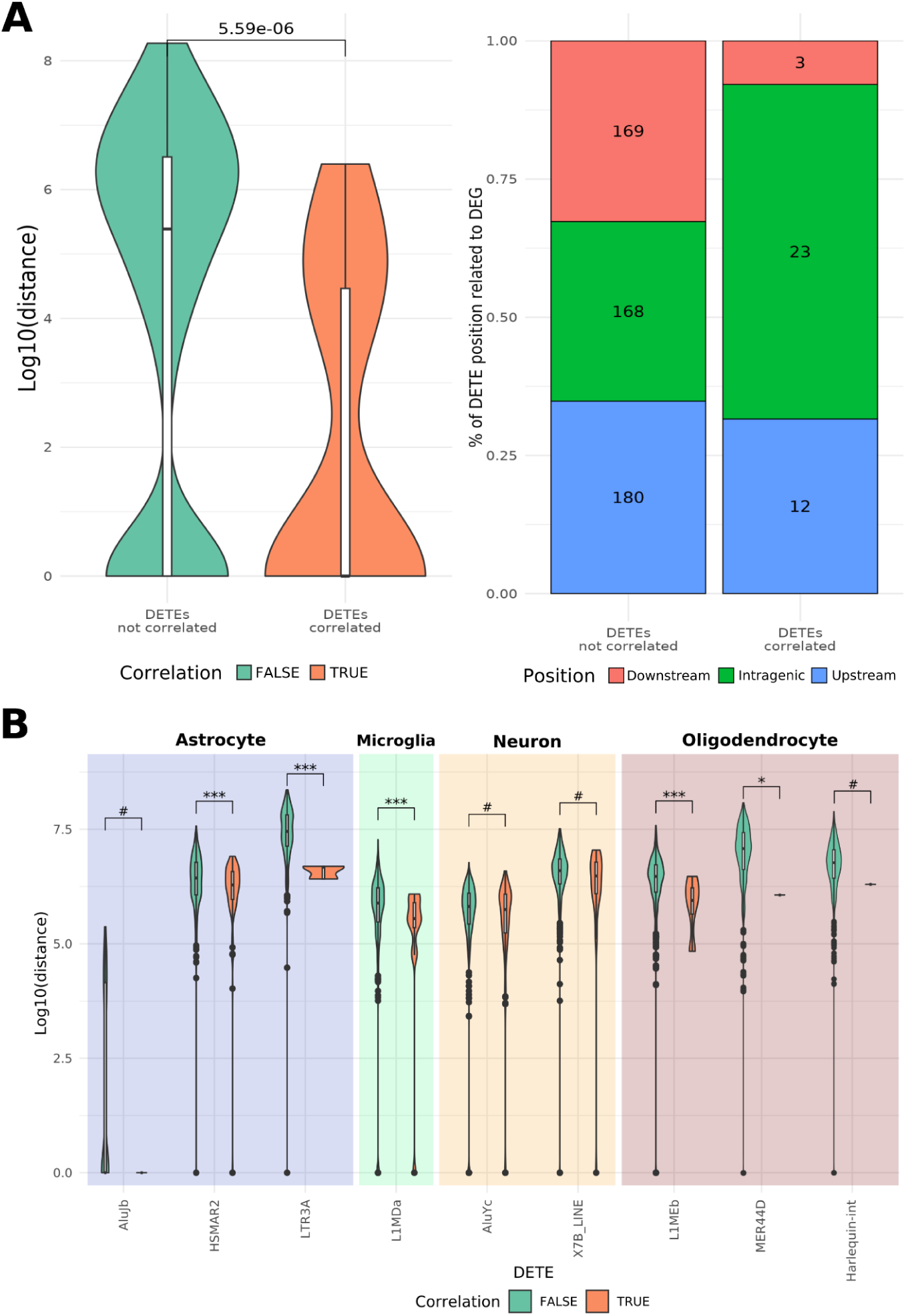
Distance distribution comparisons between DEGs and correlated and non-correlated DETEs for each cell type. **A**-Grouped distance analysis in microglia. (Right) Violin plot of distance distribution of DETE to DEG distance, grouped by sex. On top of the plot, p-value of the WNW test. (Left) Barplot with the proportion and total number of types of distances comprising each group. The distance type is defined by the relative position of the DETE related to the DEG. **B**-Significant results of individual contrast of distances for each DETE, grouped by cell type (*: FDR < 0.05, **: FDR < 0.01, ***: FDR < 0.001, #: FDR < 0.1 & p < 0.05). DETE: differentially expressed transposable element, DEG: differentially expressed gene.

When performing this analysis at the subfamily level, we decided to group the distances of the five locations closest to each DEG, since using only one distance resulted in sample sizes of correlated DETEs too small to obtain statistically significant results (see Methods). When disaggregated by subfamily, although not all of them individually have a significant difference between distances (see Table S1), some of them still show a shorter distance between correlated DEGs than between non-correlated DEGs (Figure 3B).

Additionally, from these inferred associations, we constructed TE–gene networks to visualize and explore the global connectivity of TEs in the transcriptomic context of Parkinson’s disease. To further prioritize candidate regulatory relationships, associations involving TEs located within 10 kb of a gene were highlighted as potential cis-regulatory candidates, as shorter physical distances may increase the likelihood of local regulatory effects, such as TE-derived enhancers or influences on chromatin structure. Nevertheless, longer genomic distances do not exclude functional associations, since TEs can also generate regulatory RNAs capable of modulating gene expression through trans-acting mechanisms, a phenomenon previously described in multiple cellular contexts (Kojima S, 2023).

The astrocyte network comprised 146 associations, linking 125 DEGs with 8 DETE subfamilies (Figure 2A). The L1 and Alu families were the most abundant, while the DNA transposon HSMAR2 showed the highest number of predicted associations (56 DEGs), acting as a major hub. The most connected DETEs were upregulated in males, indicating potential sex-dependent TE regulatory activity. Enriched biological processes included pathways related to telomere organization and maintenance (GO:0007004, GO:0051973), whose biology involves repetitive regions of the genome, such as is the case with many TEs. Additional enriched terms included innate immune responses (GO:0002218, GO:0050778), and axon extension (GO:0045773), both highly relevant in the development of PD in this cell type (PMID: <u>35052674</u>, https://doi.org/10.1007/978-3-031-64839-7_13).

Neurons displayed the smallest and least connected network, comprising 80 associations between 77 DEGs and 8 DETE subfamilies (Figure S8A). Most DETEs were increased in females, except for LINE X7B, which showed the opposite tendency. The hAT-Tip100 and Alu families were the most frequent, and the AluYc subfamily had the highest number of predicted associations (37 DEGs). Despite its smaller size, the neuronal network was strongly associated with processes linked to Aβ peptide processing (***GO:0090083, GO:1902430***, GO:0042987), a key mechanism in neurodegeneration. Additional enriched pathways included ubiquitination and apoptotic processes (GO:2001238, GO:0070534, GO:0006513), consistent with neuronal susceptibility to proteotoxic stress, and the NF-κB signaling pathway (GO:0043123), which have been related both to neuronal survival and apoptosis in PD, depending on other processes such as neuroinflammation (<u>10.1111/ejn.15242</u>, PMC3115056) (Figure 2B).

The microglial network contained 126 associations, with the MIR family (SINE class) as the main contributor (MIRc and MIR3), together with the LINE subfamily L1MDa, all upregulated in males (Figure S8B). MIR3 was the most connected subfamily, with 58 associated DEGs, and was also linked to L1MDa through three genes: LAPTM5, MAPK1IP1L, and RNF149. Enrichment analysis revealed strong associations with pro-inflammatory cytokine production pathways (GO:0032728, ***GO:1900017***, GO:0001816). Key genes included LAPTM5 and FLOT1, whose predicted associations were supported by close genomic proximity to MIR3. Other examples were CD300A and PCSK5, which contained intragenic MIRc sequences, and IL17RA, previously reported as essential in neuroinflammation and neurodegeneration (PMID: 31351185). Additional enriched pathways included the p53 pathway (GO:1901798) and the MAPK cascade (GO:0000165), both relevant to microglial activation and function (Figure 2C).

The oligodendrocytes displayed the largest association network, with 658 associations involving 287 DEGs and 49 DETE subfamilies. These include the LTR, DNA, and LINE classes (Figure S8C), with the ERV1, L1, and TcMar-Tigger families emerging as the most frequent participants. L1ME5 and HAL1M8 subfamilies, both from the L1 family, showed the highest number of predicted associations (44 and 41 DEGs respectively), were upregulated in females, and shared many associated genes. Most genes linked to these DETEs were enriched in DNA metabolic and processing pathways (GO:0006259). Enrichment of all genes in the network also revealed terms related to gene regulation and transcription (***GO:0006357, GO:0016575***), as well as viral genome replication (***GO:0039694***), which is biologically consistent with TE biology, as both processes involve genomic insertion events (PMID: <u>35228718</u>). Additional enriched processes of interest included myelination (***GO:0120035***, GO:0022010) and oxidative stress responses (GO:0034599), which are closely connected to oligodendrocyte function and Parkinson’s disease pathology (Peferoen et al., 2014, PMCID: <u>PMC9303520</u>).

### Dysregulated TEs are closer to PD hallmark genes, mainly in neurons

Apart from sex differences, TE expression dysregulation and its mechanisms involved in PD remain widely unexplored, with existing information only available at the family level. To delve deeper into this field, specifically into the possible relationship between transcriptional dysregulation of TEs and PD pathology, we decided to investigate the TE-distance landscape of PD hallmark genes. This was done by comparing the closest distances between a panel of the 60 highest-ranking genes associated with PD and DETEs and non-significant ones, at the TE subfamily level and by cell type. Overall, all cell types except oligodendrocytes showed that their DETEs were significantly closer to most PD genes (Figure 4A). Notably, neurons obtained significant results across the entire gene panel and a larger effect size in all cases, meaning a greater difference between the distances of DETEs and non-significant ones (all results in Table S2). Interestingly, we found core PD genes (known as PARK genes (<u>Full table</u>)) among the genes with the highest significance and effect size, whose distributions also showed a clear tendency to have DETEs closer to their genomic position, as is the case with SNCA (PARK1/4) and DJ-1 (PARK7) in neurons, PINK1 (PARK6) in astrocytes or GBA1 (considered one of the most significant genetic risk factor for PD [PMID: 30702840, PMID: 39267121]) in microglia, among others (Figure 4B).

**Figure 4.**
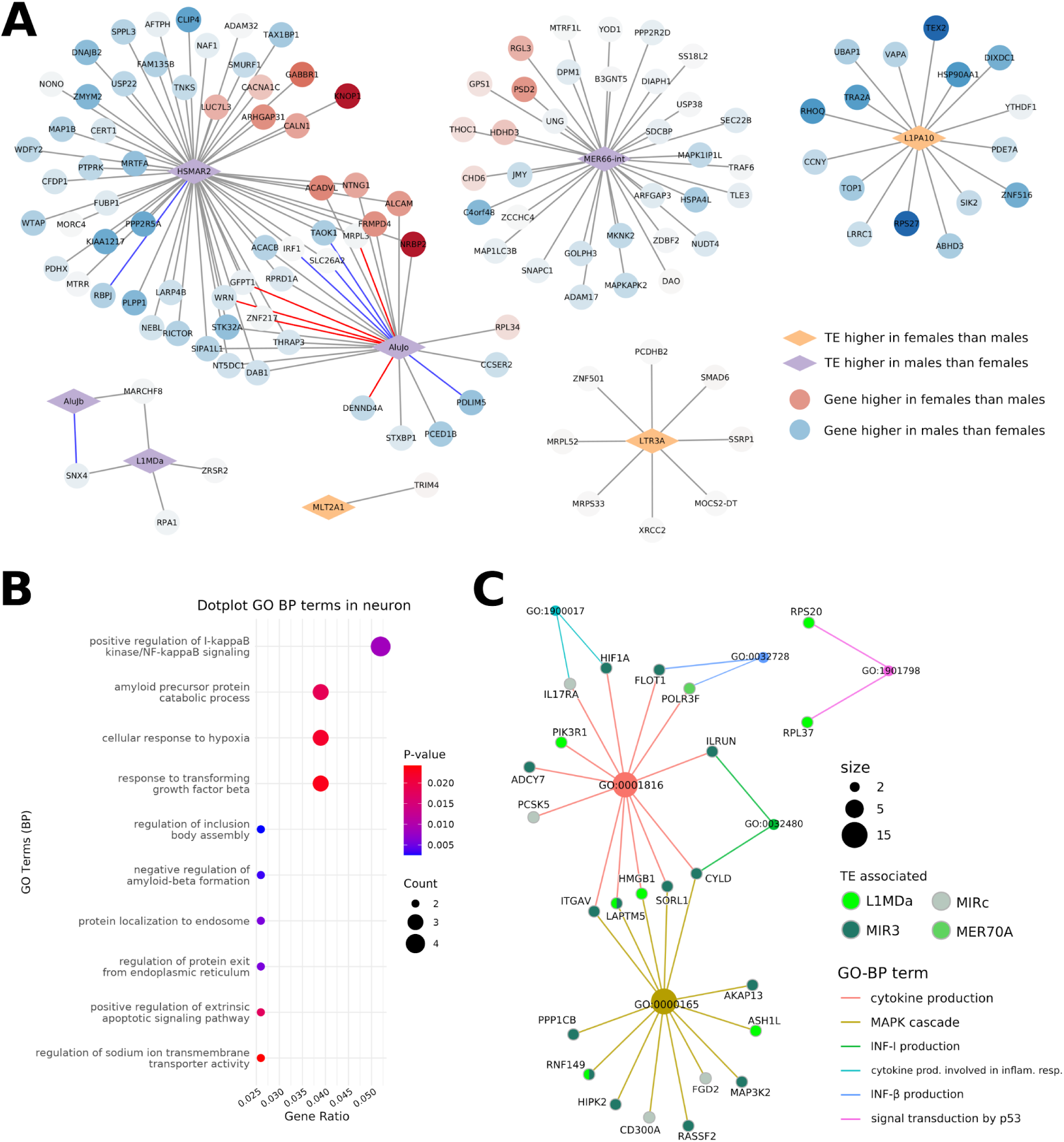
DETE-gene association map and possible biological implications of their dysregulation. **A-**Correlation network between DETEs and DEGs detected in astrocytes. associations in red are those that, in addition to having a significant logFC correlation, are also supported by a short genomic distance between the nodes (<10 kb). **B-** Dotplot of most relevant GO biological processes overrepresented in DEGs significantly correlated with DETEs in neuronal network, ordered by p-value. **C-** CNplot of inflammation related processes enriched in microglia network. DEGs nodes are coloured by the DETE to which they are correlated to. Colour of the edges represent the biological process related to the connected nodes.

**Figure 5.**
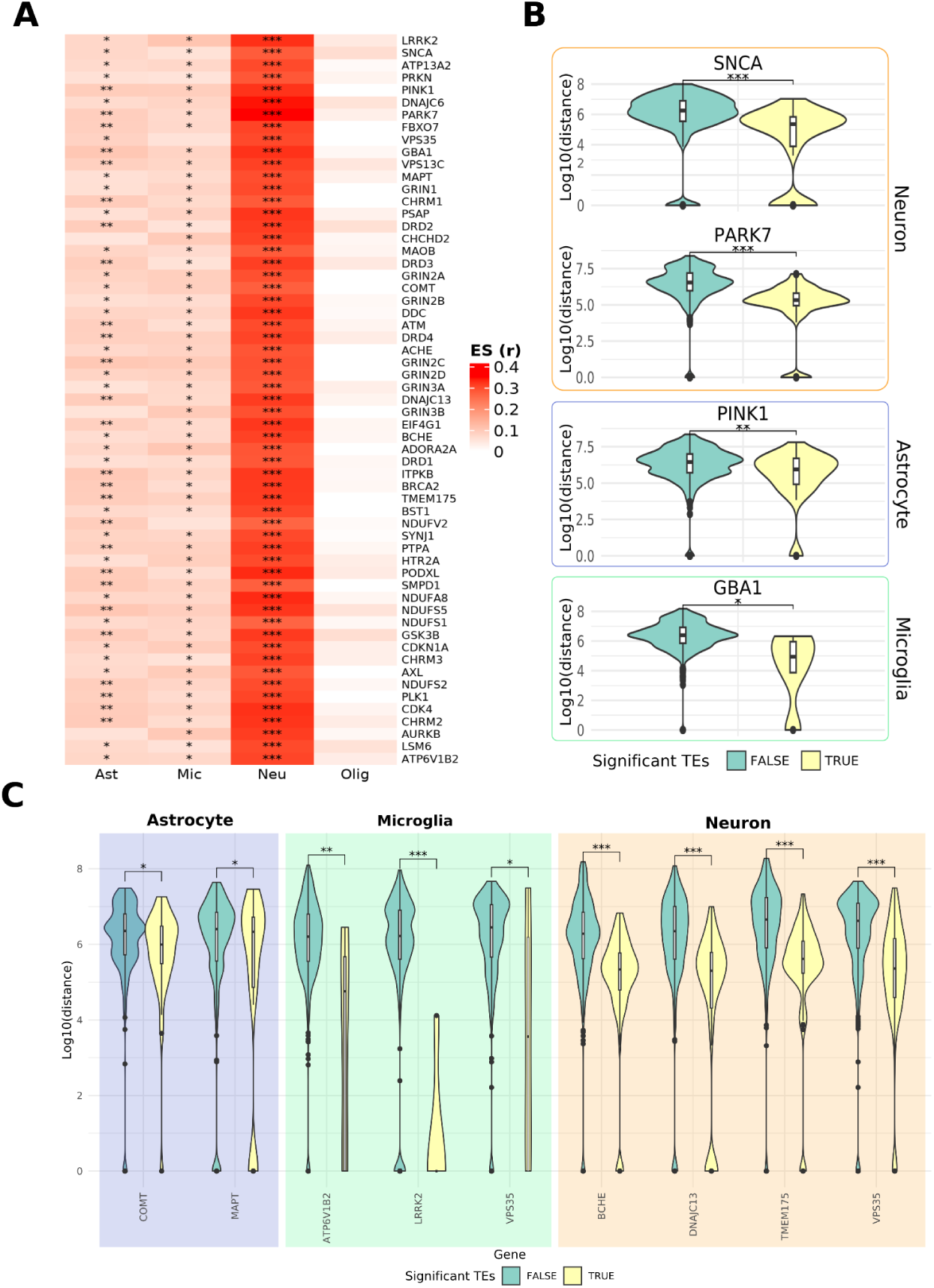
Distance contrast between DETEs and non-significant TEs in PD hallmark genes. **A**-Heatmap with effect size and significance of distance contrast in top 60 PD-related genes, divided by cell type. **B**-Violin plot of distance values (in log10(bp)) between DETEs (yellow) and non-significant TEs (green) in different PD-core genes. **C**-Violin plot of distances distribution in DETEs (yellow) and non-significant TEs (green) from genes that were both in the top 60 PD-related genes and differentially expressed in PDvsC contrast in each cell type. ES (r): Effect Size (Wilcoxon r), bp: base pair, Ast: Astrocytes, Mic: Microglia, Neu: Neurons, Oli: Oligodendrocytes, *: FDR < 0.05, **: FDR < 0.01, ***: FDR < 0.001.

For a more cell-specific view, we decided to select from the initial pool of PD hallmark genes those that were also significant in our DEA for the PDvsC contrast, for each cell type. This allowed us to identify those genes that are differentially expressed in a cell-type-specific manner and are significantly closer to their DETEs, thus suggesting PD-related processes by cell type that may be influenced by the transcriptional activity of TEs (Figure 4C).

In microglia, we identified LRRK2 (PARK8), VPS35 (PARK17), and ATP6V1B2 genes, all with mutations highly related to PD development and mechanisms of action that suggest associations between them (<u>/10.1038/s42003-025-08544-4</u>, PMID: <u>41164908</u>, PMID: <u>28960461</u>). These genes are broadly related to vesicle trafficking, phagocytosis, and, generally, how effectively microglia process phagocytosed material, which are processes that fail in PD-affected microglia, leading to the accumulation of aggregates and harmful neuroinflammation (<u>10.1016/j.arr.2024.102373</u>, <u>10.1126/sciadv.adj1205</u>). In astrocytes, we identified the catechol-O-methyltransferase (COMT) gene, related to the metabolism of dopamine or L-DOPA, and the tau protein (MAPT) gene, related to microtubule stabilization, possibly indicating a shift from a homeostatic to a reactive state caused by a catechol oxidative load and aggregated protein burden (PMCID: <u>PMC10703796</u>). Finally, in neurons, we identified the BCHE gene, related to cholinergic synapses and variants related to PD (), and TMEM175, DNAJC13, and VPS35, also related to vesicle processing and autophagy, whose impairment can cause defects in receptor recycling and increased alpha-synuclein aggregation, directly damaging neurons (PMCID:<u>PMC5338534</u>, PMID:<u>25118025</u>, <u>10.1093/hmg/ddy003</u>).

Overall, our meta-analysis reveals a robust, cell-type-specific landscape of TE dysregulation in PD, with neurons showing the strongest activation, particularly of L1 subfamilies. We also uncover sex-dependent TE signatures, with distinct DETEs contributing differently in males and females. By linking DETEs with DEGs, genomic proximity, and enriched pathways, we show that TEs may participate in key PD mechanisms, including neuroinflammation, proteostasis, and oxidative stress. These findings position TEs as active regulators that may help explain the cellular and sex-specific heterogeneity of PD.

## Discussion

Sporadic PD is a multifactorial health challenge where most molecular mechanisms have not been completely understood. Additionally, sex differences—which involve substantial clinical, symptomatic, and prevalence changes—remain still partially characterized (Patel & Kompoliti, 2023; Schaffner et al., 2025). In the search for novel pathogenic pathways to dissect this gap, TEs have emerged as potential key players in multiple aging-related neurodegenerative diseases, with growing evidence linking their dysregulation to PD hallmark processes such as neuroinflammation and oxidative stress (Ravel-Godreuil et al., 2021; Singh et al., 2024). However, the precise mechanisms by which TEs contribute to the onset of PD is not yet fully understood, and it remains unclear whether TE activation is a primary cause of neurodegeneration or a secondary consequence of other pathogenic processes. In addition, much of the existing literature on TEs in PD is limited to the TE family level or focused on specific loci within a single family, with cell-type resolution studies remaining also highly limited, leaving the activation dynamics of hundreds of TE subfamilies in different cell types still largely unexplored (Deng et al., 2024; Garza et al., 2025). A previous retrotransposons atlas in NDs has described a highly inconsistent TE expression across different studies, emphasizing the urgent necessity of applying meta-analysis techniques to extract a common and robust signal (Deng et al., 2024). To address these knowledge gaps, our study provides the first comprehensive single-cell meta-analysis atlas of TE expression in the SNpc with an additional sex specific scope, overcoming the consistency limitations of individual studies and providing a trustworthy cell-type specific signature of TE dysregulation supported by all available data to date.

Firstly, the study identified TEs with cell-type-specific expression, consistent with other single-cell studies (Deng et al., 2024). These TE markers, although less pronounced than protein-coding genes, reveal cell-type-specific dynamics of TEs, underscoring their potential relevance in specific cellular functions and differentiation fate (PMID: <u>39088190</u>)(Deng et al., 2024). Additionally, the transcriptional derepression of TEs observed between controls and PD patients in our results aligns with the hypothesis that aging and neurodegeneration are associated with an heterochromatin instability that triggers the reawakening of these dormant elements (Copley & Shorter, 2023; Ravel-Godreuil et al., 2021). Our findings reveal that neurons undergo the most extensive TE activation, specifically within the L1 family, where 39 subfamilies were significantly upregulated in PD. This activation is likely facilitated by the loss of repressive epigenetic marks such as DNA methylation, a process that can be triggered by the high levels of reactive oxygen species (ROS) characteristic of the dopaminergic environment (Baeken et al., 2020; Bonnifet et al., 2025; Chesnokova et al., 2022; Garza et al., 2025). Our spatial analysis revealed that DETEs are significantly closer to core PARK genes than non-dysregulated TEs, particularly in neurons. The proximity of these elements to hallmark loci—such as SNCA and PARK7 in neurons, PINK1 in astrocytes, or GBA1 in microglia—suggests that TEs may act as cis-regulatory elements, potentially disrupting gene regulatory networks by acting as alternative promoters or enhancers in the human brain, and promoting their dysregulation during PD pathology (Chesnokova et al., 2022; Gordevičius et al., 2023; Mustafin, 2025).

Our analyses also identified sex-specific DETE subfamilies that have been previously linked to neurodegenerative diseases pathological processes, and have been also related to biological networks highly related to PD biology. In neurons, we observed significant activity in subfamilies such as AluYc, Alu or MER66C among others. The Alu family, particularly primate-specific subfamilies like AluYc, which emerged as the most connected hub in our neuronal network, has been increasingly recognized for its role in transcriptional noise and mitochondrial dysfunction (Baeken et al., 2020; Chesnokova et al., 2022; Fröhlich et al., 2024), and have also been related with overexpression of CNTN5, related with PD through LRRK2 (Khan et al., 2024). In microglia, although the SDID contrast did not yield any significant DETE, we identified some of them with opposite patterns in IDF and IDM that were also largely connected with important coding genes, highlighting subfamilies like L1MDa or MIR3 and MIRc from MIR SINE family. Recent evidence suggests that the activation of LINE-1 in microglia is coupled to a “viral mimicry” response related to IFN signaling cascade that sustains chronic neuroinflammation (Garza et al., 2025; Ruffo et al., 2025), and also have been reported to be related with Alzheimer’s-relevant pathways like Aβ phagocytosis or cytokine secretion (PMID: <u>39604588</u>)(Singh et al., 2024). Furthermore, the MIR family can encode regulatory RNAs that can modify the chromatin organization and re-wiring boundaries between active and repressive domains (<u>PMC4538669</u>), possibly shifting them to a more inflammatory phenotype. Thus, these subfamilies could be potentially activating pro-inflammatory signaling mainly in male microglia, in concordance with the enriched biological processes obtained in their correlated genes (Bourque et al., 2023; Patel & Kompoliti, 2023). In oligodendrocytes, the most represented family in sex-specific DETEs is the HERVs TEs, mostly increased in males and with subfamilies of interest including MLT1G3 or Harlequin-int, with highest logFC. Although HERVs have been described to be protective in astrocytes in PD (PMID: <u>39840837</u>), it has been described that HERVs expressed in glia can drive pro-inflammatory and demyelinating cascades (PMID: <u>38927681</u>), and that the ERV RNLTR12-int is crucial for myelination (<u>10.1016/j.cell.2024.01.011</u>). Thus, these HERVs subfamilies could be affecting myelination and inflammatory response specifically in male oligodendrocytes, which also appear enriched in genes from regulatory networks proposed (Peferoen et al., 2014; Talevi et al., 2025).

The biological networks characterized in this study reveal striking sex-specific differences in TE-driven pathways. In neurons, the network was predominantly upregulated in females and showed a strong negative correlation with genes related with Aβ peptide processing, ubiquitination, and apoptotic processes (PMID: <u>39604588</u>)(Singh et al., 2024). This may indicate that TE sex-differential dysregulation in neurons may be intrinsically linked to divergent clearing misfolded proteins capacity between sexes, leading to a different proteotoxic stress impact previously described (Gordillo-González et al., 2024; Patel & Kompoliti, 2023)(PMID: <u>31133790</u>). In contrast, the astrocyte network was dominated by the DNA transposon HSMAR2, which acted as a major hub upregulated in males. This male-biased network was significantly enriched for innate immune responses and telomere maintenance. The link between TEs and telomere biology is particularly relevant, as telomeric regions are composed of repetitive sequences that share maintenance mechanisms with certain TEs (Copley & Shorter, 2023; Talevi et al., 2025). Given that males generally exhibit higher levels of neuroinflammation and an earlier onset of neurodegeneration in the substantia nigra, our findings imply that male-specific TE activation may be a primary driver of this pro-inflammatory environment, which can contribute directly and indirectly to oxidative stress and neuronal degeneration (Boyd et al., 2022; Mukhara et al., 2020). Furthermore, the observation that sex hormones like estradiol and testosterone heavily influence the epigenetic machinery responsible for TE silencing—such as DNA methyltransferases (DNMTs)—buttresses the hypothesis that the sex-biased activation of these elements is a key mediator of the molecular differences in PD pathogenesis (Bourque et al., 2023; Gordillo-González et al., 2024; Terrin et al., 2023; Yanguas-Casás, 2020).

The integration of these findings suggests that TEs are not merely secondary consequences of neurodegeneration but active mediators of sex-differential biological mechanisms. The differential methylation of TEs, which is highly influenced by biological sex and hormonal status, may explain why certain pathogenic pathways are more active in one sex than the other (Hyacinthe & Bourque, 2024; Stoccoro, 2025). For example, male-specific hypomethylation at the PARK7 locus has been reported, matching our observations of male-biased TE activity in regions associated with neurotransmission guidance (Kochmanski et al., 2022). These insights open new avenues for precision medicine, where targeting specific TE activity could mitigate sex-specific pathological trajectories.

Despite the strengths of this meta-analysis, there are some challenges associated with the heterogeneity of TE that require further research before they can be overcome. The diverse mechanisms of action proposed for their pathogenicity—ranging from insertional mutagenesis and genomic instability to transcriptional toxicity and viral mimicry—makes it extremely difficult to perform a single analysis that encompasses all facets of their biology. Many of these functions depend on genomic location, methylation status, RNA binding capacity, among others. This would require a more detailed analysis of the TE subfamilies highlighted in this work, using multi-omic information that would allow a complete characterization of the causal relationship between TE activation and the initiation of PD. However, the results of this study have identified TE subfamilies whose activation could be highly relevant to the sex differences identified in PD in a cell-type-specific manner, also proposing the genes through which they exert their effect. We obtained these results through an in silico analysis of data available in public repositories, which demonstrates the power of using this type of approach to obtain more insight into the etiopathogenesis of different diseases from previously generated data. To promote these approaches, it is essential to conduct open research and make data available in public repositories according to FAIR principles (Findable, Accessible, Interoperable, Reusable), with full information on covariates of interest, such as sex. We also acknowledge that these results might differ if performed in other brain regions, since it has been described that TE expression significantly varies between tissues and, more specifically, brain regions (Bogu et al., 2019; Garza et al., 2025). However, the number of sc-RNAseq studies available from other brain regions that meet our systematic review criteria was not enough to perform a meta-analysis, thus we only analyzed the SNpc studies, which is also a central region during PD pathogenesis (Braak et al., 2003). Finally, we contributed to open science with a public webtool that offers a valuable resource for the community to further investigate TE expression patterns in PD.

## DECLARATIONS

### Ethics approval and consent to participate

Not applicable.

### Consent for publication

Not applicable.

### Availability of data and materials

An interactive webserver to explore the scRNA-seq count data is freely available at https://bioinfo.cipf.es/cbl-atlas-pd/. Source data are provided with this paper.

### Competing interests

The authors state no conflict of interest.

### Funding

This research was supported and partially funded by CIAICO/2023/149, CIACIF/2021/221 and CIGE/2024/242 funded by the Consellería de Educación, Cultura y Universidades de la Generalitat Valenciana, PID2021-124430OA-I00 funded by MCIN/AEI/10.13039/501100011033 and by “ERDF A way of making Europe”.

### Author contributions

Conceptualization: FrGG, MKI; Funding acquisition: FrGG; Data Curation: FeGG; Investigation: FrGG, MKI, HS, FeGG, MRH, CGR, RGR; Bioinformatic Analysis: FeGG, RGR, ISS; Software: FeGG, RGR; Supervision: FGG, CGR, MKI; Writing-Original Draft Preparation: FrGG, MKI, HS, FeGG, MRH, CGR, RGR, ISS. All authors read and approved the final manuscript.

## Acknowledgements

The authors thank the Principe Felipe Research Center (CIPF) for providing access to the cluster, cofunded by European Regional Development Funds (FEDER) to the Valencian Community 2014-2020. We would like to thank Dr Shohei Kojima of the Keio University Human Biology-Microbiome-Quantum Research Centre (WPI-Bio2Q), Tokyo, 160-8582, Japan, for his guidance. The authors also thank Stuart P. Atkinson for reviewing the manuscript.

## Supplementary material

**Figure S1.**
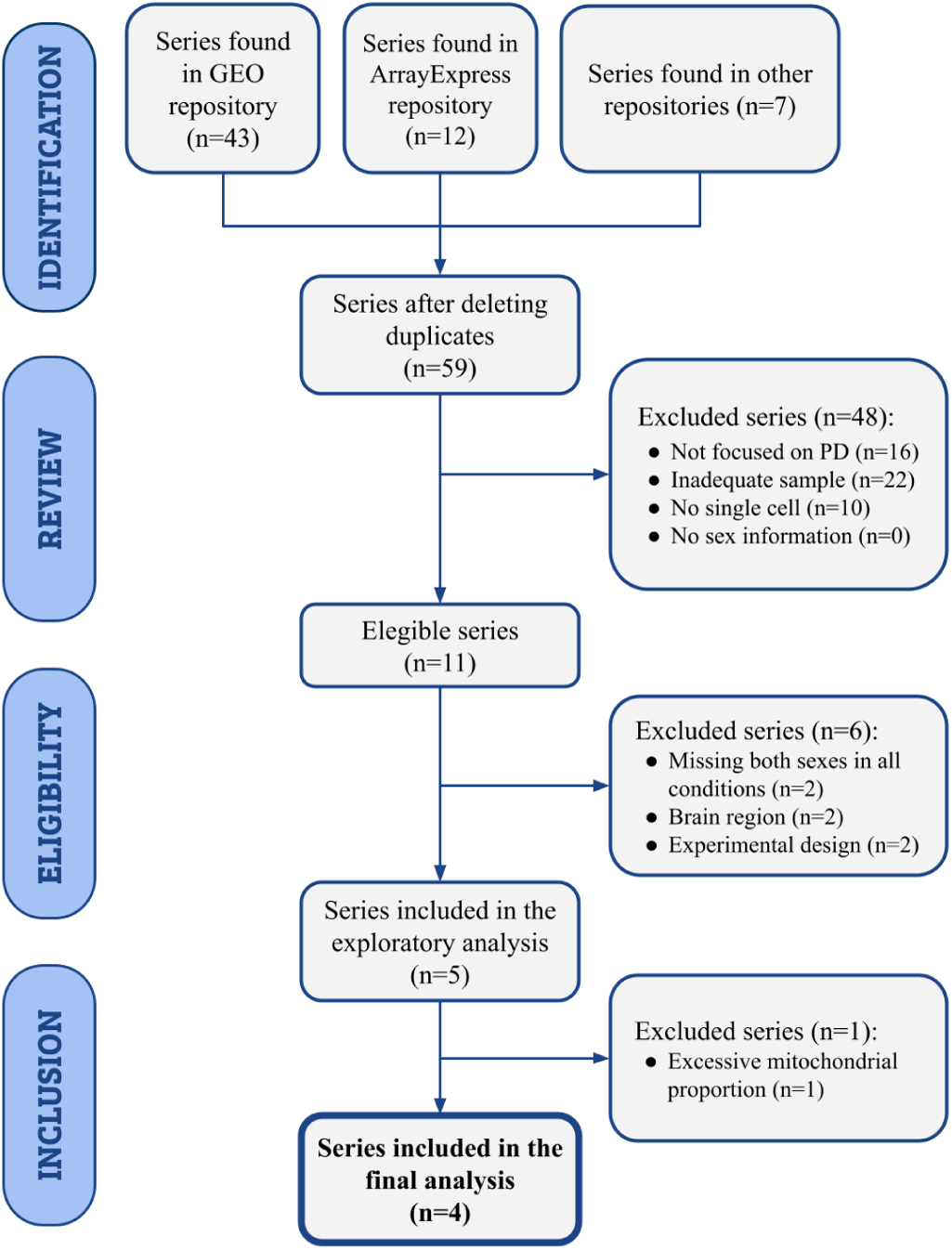
PRISMA diagram results

**Figure S2.**
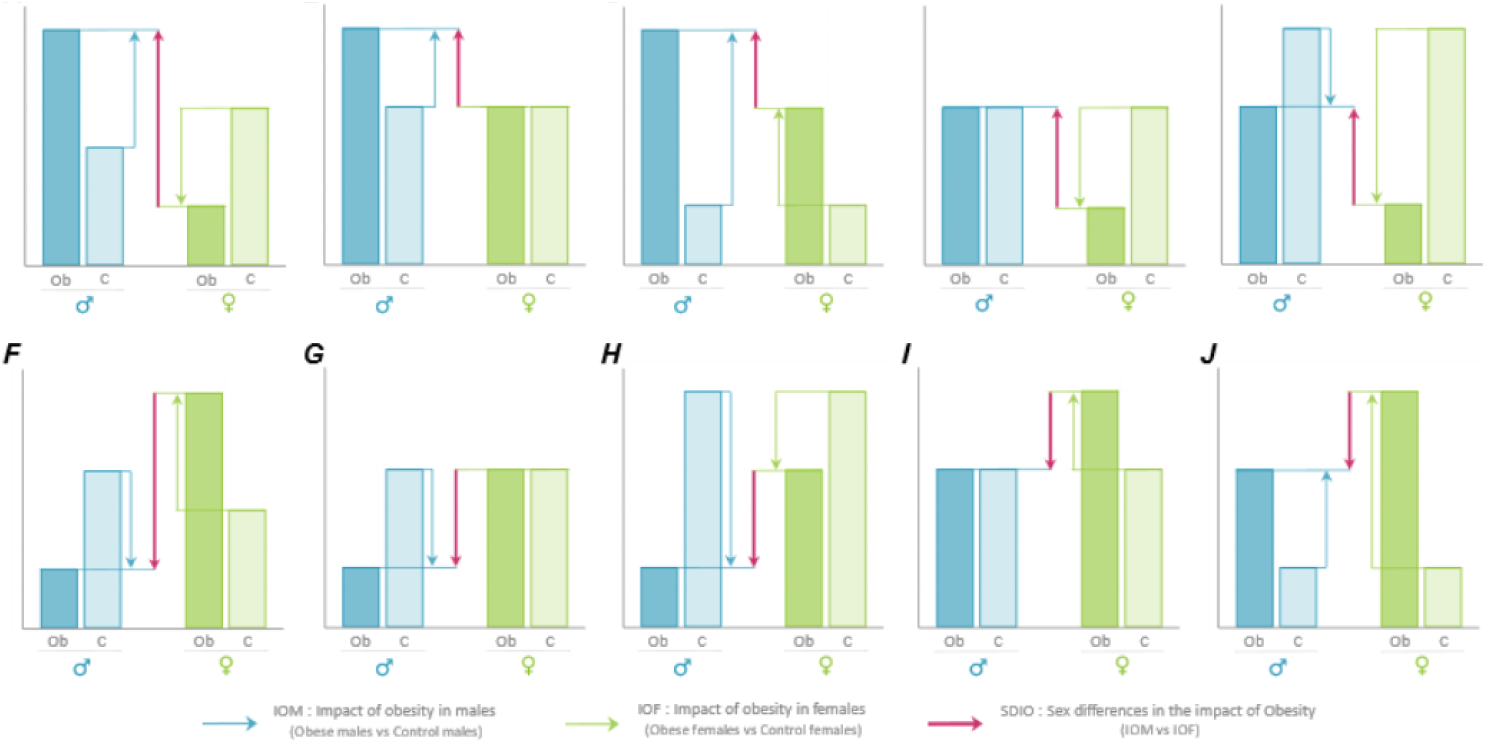
Possible scenarios of significant expression change in the Sex Differential Impact of Disease (SDID) contrast explained. The upper row (with red arrow pointing up) indicates the scenarios of a positive logFC, referred as “higher in females than male”. The lower row (with red arrow pointing down) indicates the scenarios of a negative logFC, referred as “higher in males than females”.

**Figure S3.**
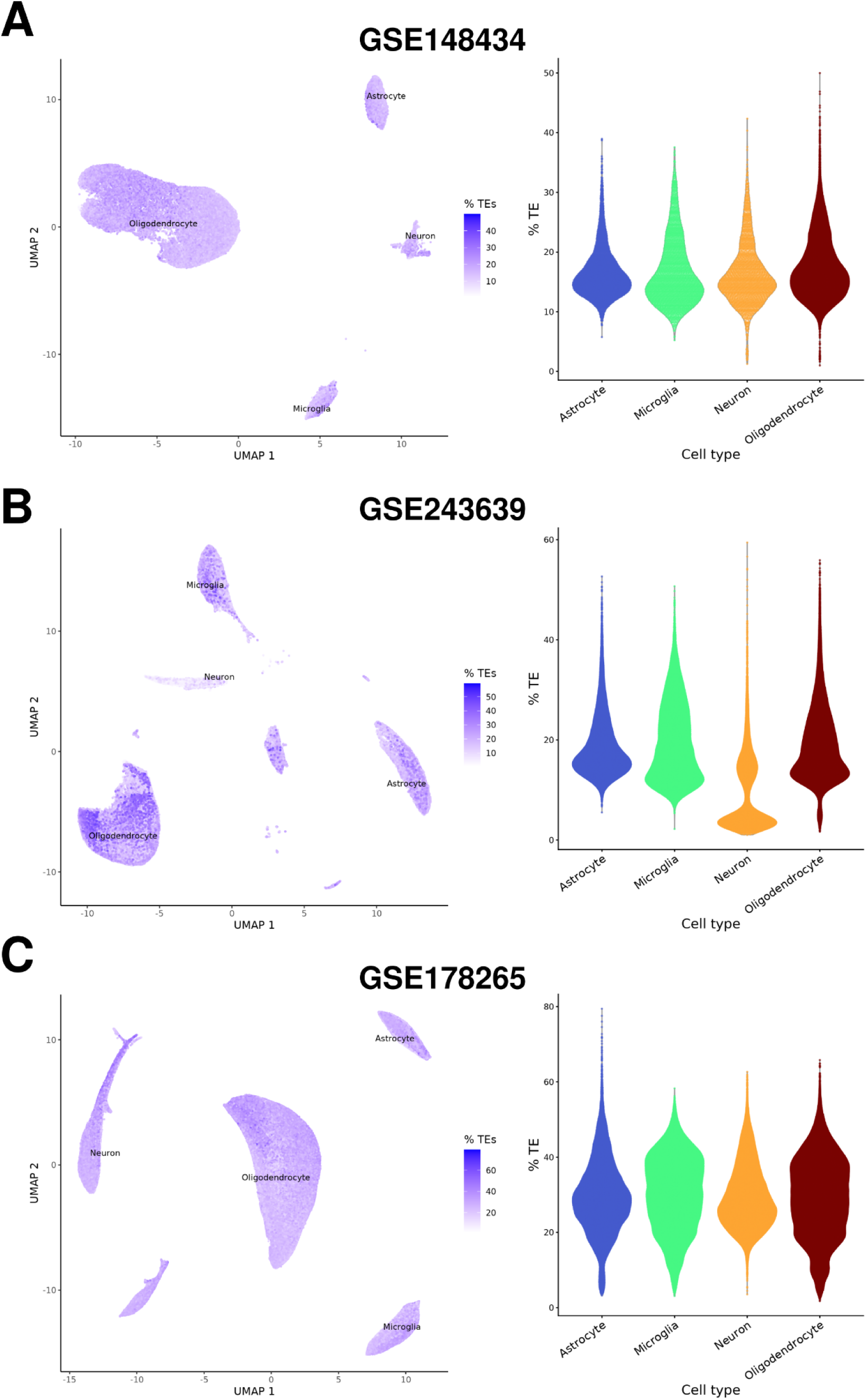
TE expression rates in other datasets.

**Figure S4.**
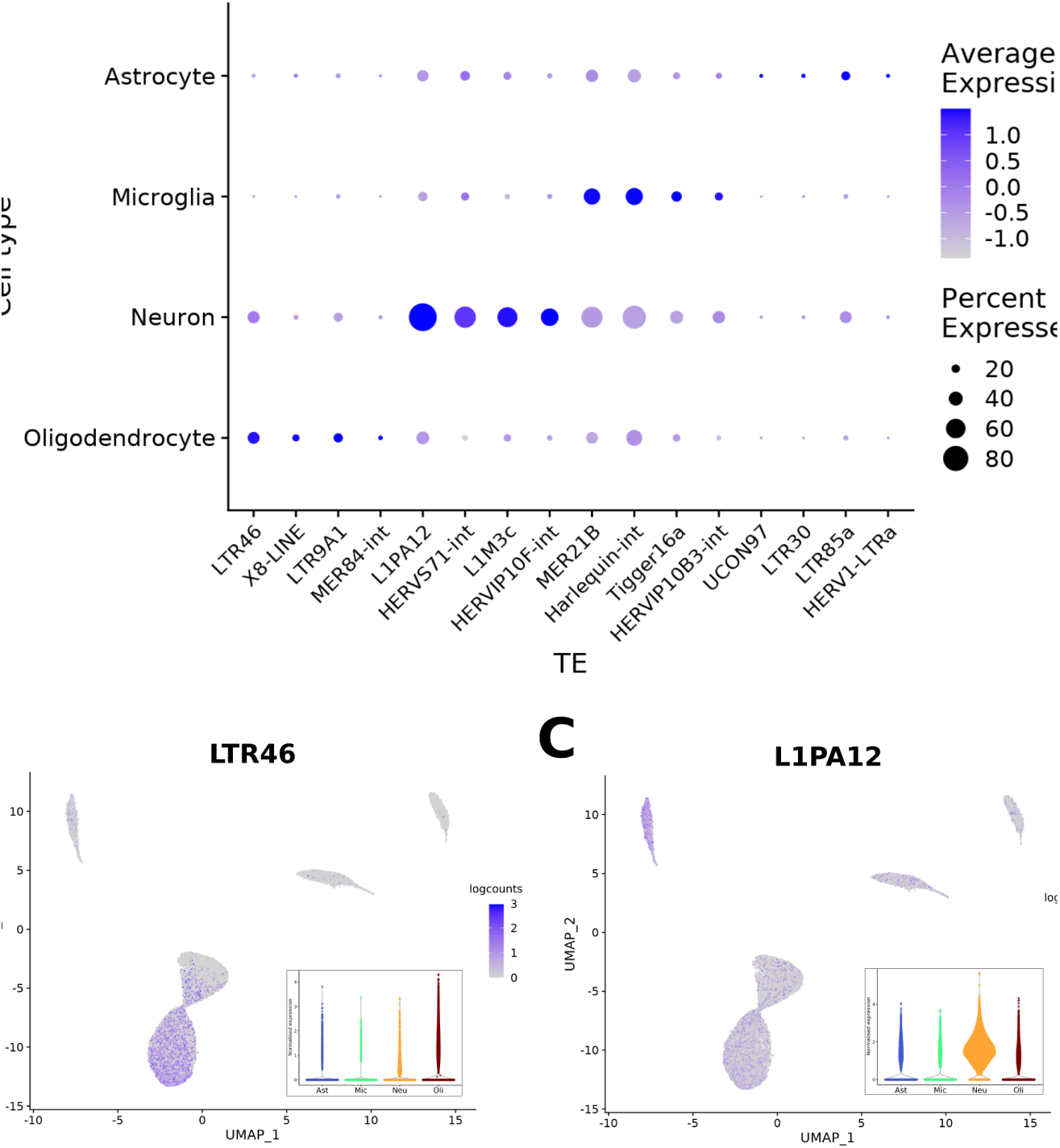
TE Markers GSE157783.

**Figure S5.**
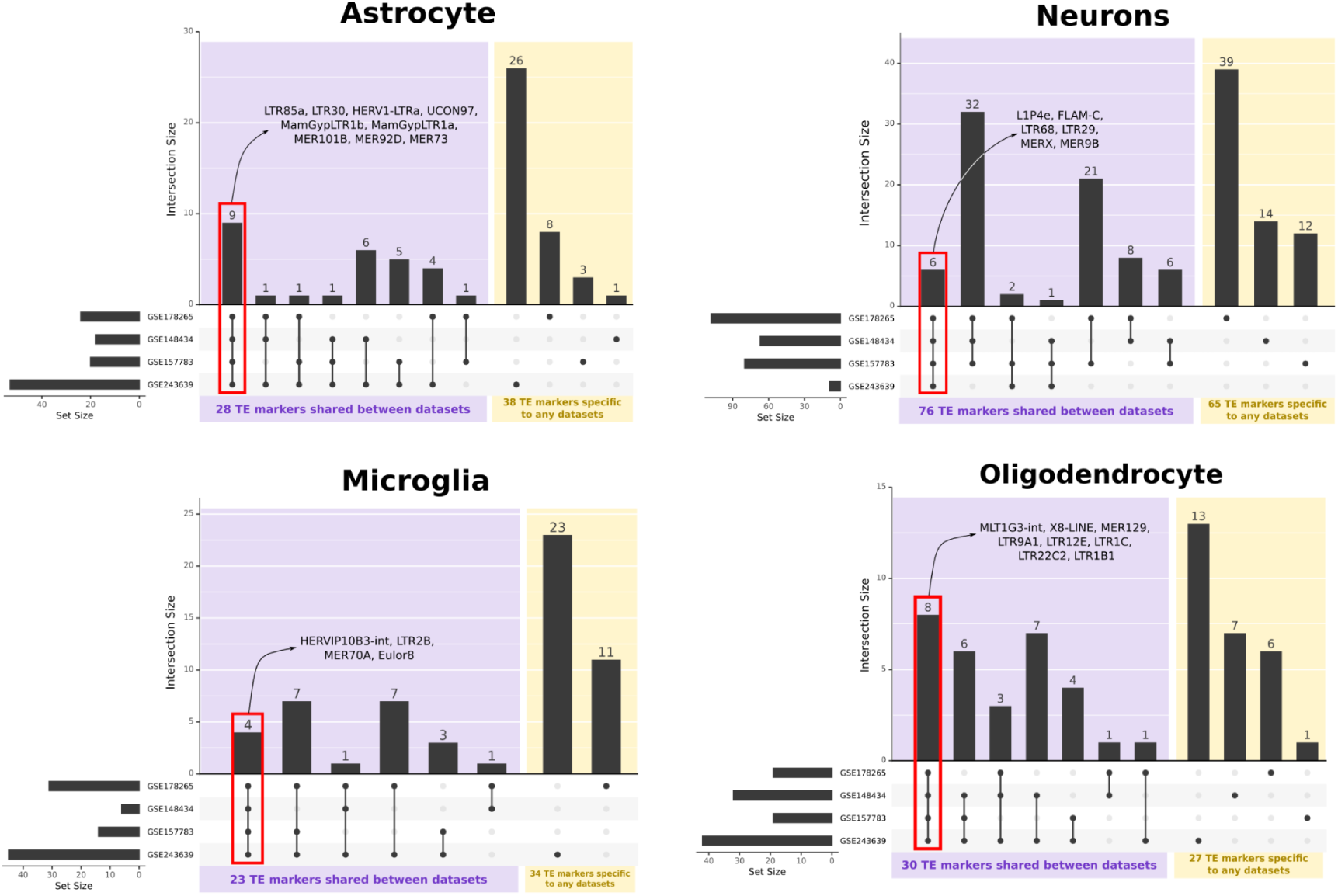
TE Markers commons between studies.

**Figure S6.**
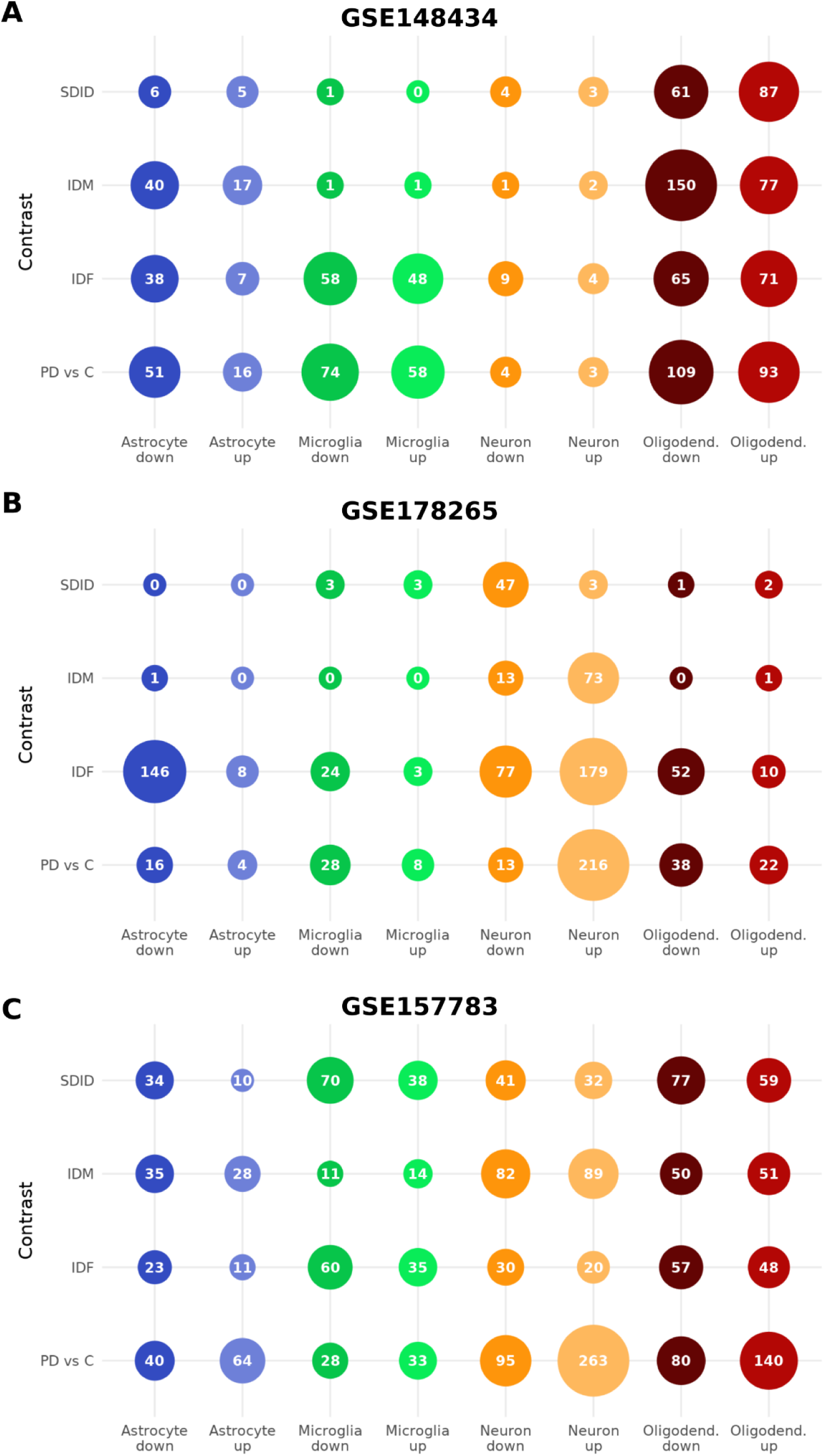
Individual studies DETEs per contrast.

**Figure S7.**
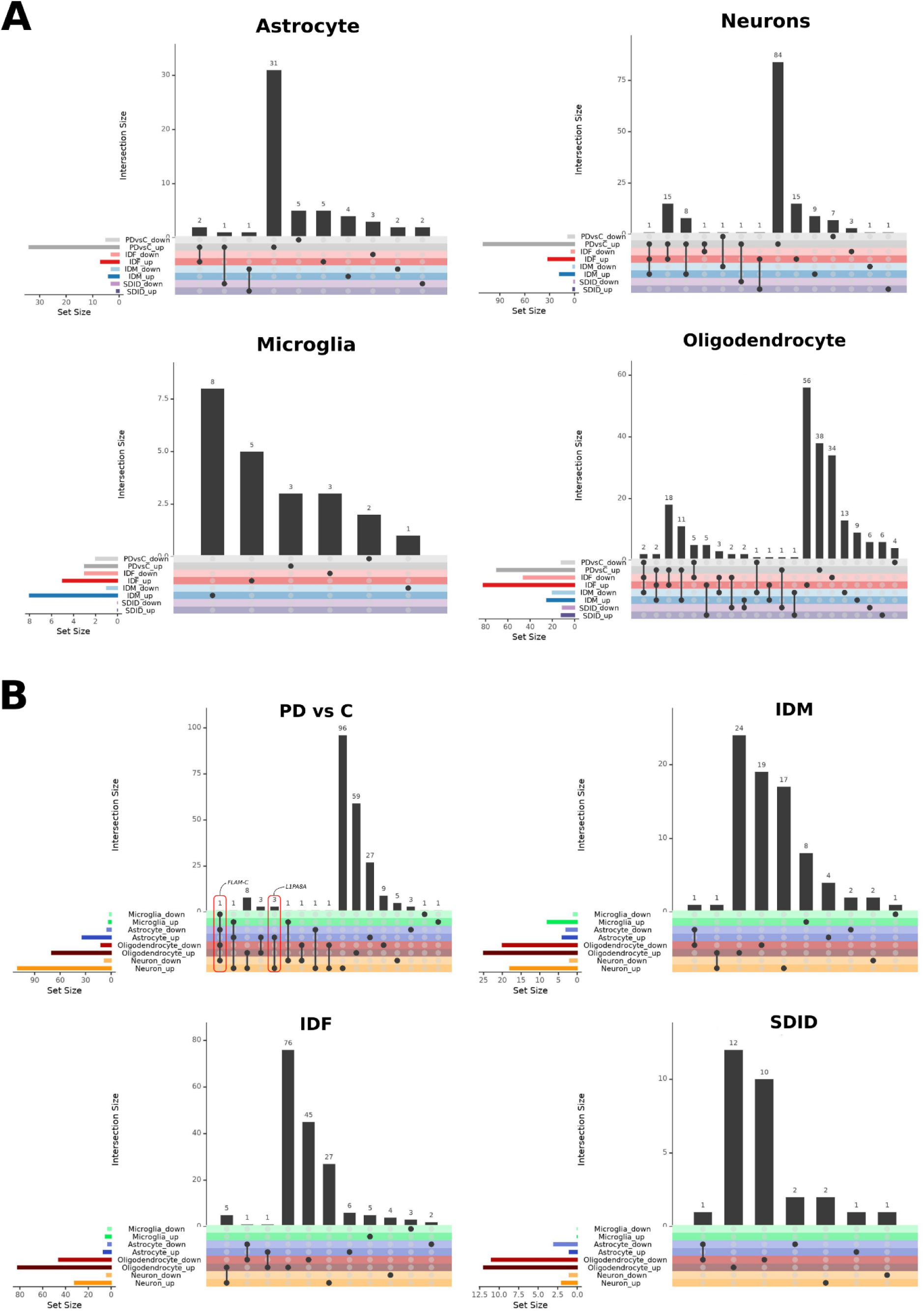
Meta-analyses upsets.

**Figure S8.**
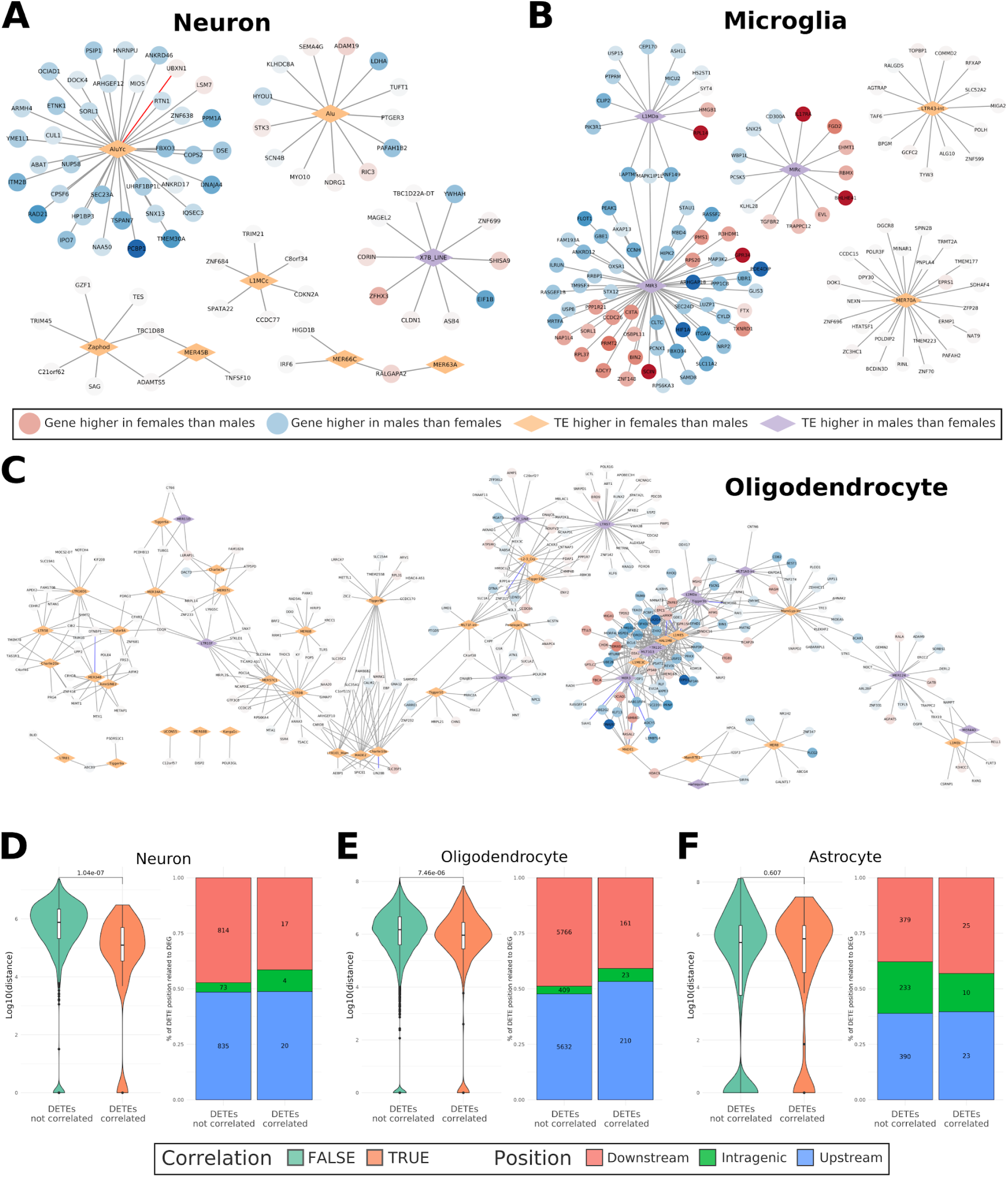
Networks and distances contrasts of other cell types.

## Notes

https://bioinfo.cipf.es/cbl-atlas-pd/

### Competing Interest Statement

The authors have declared no competing interest.

## Bibliography

Baeken, M. W., Moosmann, B., & Hajieva, P. (2020). Retrotransposon activation by distressed mitochondria in neurons. Biochemical and Biophysical Research Communications, 525(3), 570–575. 10.1016/j.bbrc.2020.02.106

Bogu, G. K., Reverter, F., Marti-Renom, M. A., Snyder, M. P., & Guigó, R. (2019). Atlas of transcriptionally active transposable elements in human adult tissues (p. 714212). bioRxiv. 10.1101/714212

Bonnifet, T., Sinnassamy, S., Massiani-Beaudoin, O., Mailly, P., Monnet, H., Loew, D., Lombard, B., Servant, N., Joshi, R. L., & Fuchs, J. (2025). Steady-state neuron-predominant LINE-1 encoded ORF1p protein and LINE-1 RNA increase with aging in the mouse and human brain. eLife, 13, RP100687. 10.7554/eLife.100687

Bourque, M., Morissette, M., Soulet, D., & Di Paolo, T. (2023). Impact of sex on neuroimmune contributions to Parkinson’s disease. Brain Research Bulletin, 199, 110668. 10.1016/j.brainresbull.2023.110668

Boyd, R. J., Avramopoulos, D., Jantzie, L. L., & McCallion, A. S. (2022). Neuroinflammation represents a common theme amongst genetic and environmental risk factors for Alzheimer and Parkinson diseases. Journal of Neuroinflammation, 19(1), 223. 10.1186/s12974-022-02584-x

Braak, H., Tredici, K. D., Rüb, U., de Vos, R. A. I., Jansen Steur, E. N. H., & Braak, E. (2003). Staging of brain pathology related to sporadic Parkinson’s disease. Neurobiology of Aging, 24(2), 197–211. 10.1016/S0197-4580(02)00065-9

Chesnokova, E., Beletskiy, A., Kolosov, P., Chesnokova, E., Beletskiy, A., & Kolosov, P. (2022). The Role of Transposable Elements of the Human Genome in Neuronal Function and Pathology. International Journal of Molecular Sciences, 23(10). 10.3390/ijms23105847

Copley, K. E., & Shorter, J. (2023). Repetitive elements in aging and neurodegeneration. Trends in Genetics: TIG, 39(5), 381–400. 10.1016/j.tig.2023.02.008

Deng, W., Citu, C., Liu, A., & Zhao, Z. (2024). Dynamic dysregulation of retrotransposons in neurodegenerative diseases at the single-cell level. Genome Research, 34(10), 1687–1699. 10.1101/gr.279363.124

Fröhlich, A., Pfaff, A. L., Middlehurst, B., Hughes, L. S., Bubb, V. J., Quinn, J. P., & Koks, S. (2024). Deciphering the role of a SINE-VNTR-Alu retrotransposon polymorphism as a biomarker of Parkinson’s disease progression. Scientific Reports, 14(1), 10932. 10.1038/s41598-024-61753-5

Garza, R., Adami, A., Thiruvalluvan, A., Wijesinghe, S., Curle, A., Tam, O., Forcier, T., Lagka, D. A., Kazakou, N. L., Atacho, D. A. M., Sharma, Y., Jönsson, M., Horvath, V., Bermudez, S., Johansson, J., Rainbow, D. B., Castilla-Vallmanya, L., Jones, J. L., Quaegebeur, A.,…Jakobsson, J. (2025). Activation of transposable elements is linked to a region- and cell-type-specific interferon response in Parkinson’s disease (p. 2025.09.03.673956). bioRxiv. 10.1101/2025.09.03.673956

Gordevičius, J., Goralski, T., Bergsma, A., Parham, A., Kuhn, E., Meyerdirk, L., McDonald, M., Milčiūtė, M., Putten, E. V., Marshall, L., Brundin, P., Brundin, L., Labrie, V., Henderson, M., & Pospisilik, J. A. (2023). Human Endogenous Retrovirus Expression is Dynamically Regulated in Parkinson’s Disease (p. 2023.11.03.565438). bioRxiv. 10.1101/2023.11.03.565438

Gordillo-González, F., Soler-Sáez, I., Galiana-Roselló, C., Hidalgo, M. R., Gómez-Cabañes, B., Grillo-Risco, R., Dolader-Rabinad, B., Díez, N. del R., Virués-Morales, A., Yanguas-Casás, N., Ferrer, F. C., & García-García, F. (2024). Uncovering sex differences in Parkinson’s Disease through metaanalysis of single cell transcriptomic studies (p. 2024.12.20.628852). bioRxiv. 10.1101/2024.12.20.628852

Hyacinthe, J., & Bourque, G. (2024). Transposable elements impact the regulatory landscape through cell type specific epigenomic associations (p. 2024.08.07.606967). bioRxiv. 10.1101/2024.08.07.606967

Khan, S. S., Jaimon, E., Lin, Y.-E., Nikoloff, J., Tonelli, F., Alessi, D. R., & Pfeffer, S. R. (2024). Loss of primary cilia and dopaminergic neuroprotection in pathogenic LRRK2-driven and idiopathic Parkinson’s disease. Proceedings of the National Academy of Sciences of the United States of America, 121(32), e2402206121. 10.1073/pnas.2402206121

Kochmanski, J., Kuhn, N. C., & Bernstein, A. I. (2022). Parkinson’s disease-associated, sex-specific changes in DNA methylation at PARK7 (DJ-1), SLC17A6 (VGLUT2), PTPRN2 (IA-2β), and NR4A2 (NURR1) in cortical neurons. NPJ Parkinson’s Disease, 8, 120. 10.1038/s41531-022-00355-2

Mukhara, D., Oh, U., & Neigh, G. N. (2020). Neuroinflammation. Handbook of Clinical Neurology, 175, 235–259. 10.1016/B978-0-444-64123-6.00017-5

Mustafin, R. N. (2025). The role of retroelements in Parkinson’s disease development. Vavilov Journal of Genetics and Breeding, 29(2), 290–300. 10.18699/vjgb-25-32

Patel, R., & Kompoliti, K. (2023). Sex and Gender Differences in Parkinson’s Disease. Neurologic Clinics, 41(2), 371–379. 10.1016/j.ncl.2022.12.001

Peferoen, L., Kipp, M., Valk, P., Noort, J. M., & Amor, S. (2014). Oligodendrocyte-microglia cross-talk in the central nervous system. Immunology, 141(3), 302–313. 10.1111/imm.12163

Ravel-Godreuil, C., Znaidi, R., Bonnifet, T., Joshi, R. L., & Fuchs, J. (2021). Transposable elements as new players in neurodegenerative diseases. FEBS Letters, 595(22), 2733–2755. 10.1002/1873-3468.14205

Ruffo, P., Perrone, B., Perrone, F., Amicis, F. D., Iuliano, R., Bucci, C., Messina, A., & Conforti, F. L. (2025). The Other Side of the Same Coin: Beyond the Coding Region in Amyotrophic Lateral Sclerosis. Pharmaceuticals, 18(10). 10.3390/ph18101573

Schaffner, S. L., Tosefsky, K. N., Inskter, A. M., Appel-Cresswell, S., & Schulze-Hentrich, J. M. (2025). Sex and gender differences in the molecular etiology of Parkinson’s disease: Considerations for study design and data analysis. Biology of Sex Differences, 16(1), 7. 10.1186/s13293-025-00692-w

Singh, S., Borkar, M. R., & Bhatt, L. K. (2024). Transposable Elements: Emerging Therapeutic Targets in Neurodegenerative Diseases. Neurotoxicity Research, 42(1), 9. 10.1007/s12640-024-00688-1

Stoccoro, A. (2025). Epigenetic Mechanisms Underlying Sex Differences in Neurodegenerative Diseases. Biology, 14(1), 98. 10.3390/biology14010098

Talevi, V., Lee, H.-M., Liu, D., Beyer, M. D., Salomoni, P., Breteler, M. M. B., & Aziz, N. A. (2025). Age and Sex Effects on Blood Retrotransposable Element Expression Levels: Findings From the Population-Based Rhineland Study. Aging Cell, 24(8), e70092. 10.1111/acel.70092

Terrin, F., Tesoriere, A., Plotegher, N., & Dalla Valle, L. (2023). Sex and Brain: The Role of Sex Chromosomes and Hormones in Brain Development and Parkinson’s Disease. Cells, 12(11), 1486. 10.3390/cells12111486

Yanguas-Casás, N. (2020). Physiological sex differences in microglia and their relevance in neurological disorders. Neuroimmunology and Neuroinflammation, 7(1), 13–22. 10.20517/2347-8659.2019.31

